# PathOmCLIP: Connecting tumor histology with spatial gene expression via locally enhanced contrastive learning of Pathology and Single-cell foundation model

**DOI:** 10.1101/2024.12.10.627865

**Authors:** Yongju Lee, Xinhao Liu, Minsheng Hao, Tianyu Liu, Aviv Regev

## Abstract

Tumor morphological features from histology images are a cornerstone of clinical pathology, diagnostic biomarkers, and basic cancer biology research. Spatial transcriptomics, which provides spatially resolved gene expression profiles overlaid on histology images, offers a unique opportunity to integrate morphological and expression features, thereby deepening our understanding of tumor biology. However, spatial transcriptomics experiments with patient samples in either clinical trials or clinical care are costly and challenging, whereas histology images are generated routinely and available for many legacy prospective cohorts of disease progression and outcomes in well-annotated cohorts. Inferring spatial transcriptomics profiles computationally from these histology images would significantly expand our understanding of tumor biology, but paired data for training multi-modal spatial-histology models remains limited. Here, we tackle this challenge by incorporating performant foundation models pre-trained on massive datasets of pathology images and single-cell RNA-Seq, respectively, which provide useful embeddings to underpin multi-modal models. To this end, we developed PathOmCLIP, a model trained with contrastive loss to create a joint-embedding space between a histopathology foundation model and a single-cell RNA-seq foundation model. We incorporate a set transformer to gather localized neighborhood tumor architecture following contrastive training, which further enhances performance and is necessary to obtain robust results. We validate PathOmCLIP across five tumor types and achieve significant performance improvements in gene expression prediction tasks over other methods. PathOmCLIP can be applied to many archived histology images, unlocking valuable clinical information and facilitating new biomarker discoveries.

## 1 Introduction

Tumor morphological features derived from pathology images have long been essential for diagnostics and research on the tumor microenvironment. In recent decades, genomic information at the DNA and RNA level has been combined with histopathology images [12] to enhance our understanding of tumor biology, facilitate tumor stratification, and guide treatment strategies. Most recently, spatially-resolved transcriptomics [39, 44, 46] preserves the positional information of expression profiles within tissues up to single-cell resolution, thus overlaying expression profiles on pathology images. Spatial transcriptomics helps understand cell-to-cell interactions and multi-cellular neighborhoods and relate them to both morphological changes and tumor progression [7, 12, 17, 19, 33].

Concomitantly, whole slide image scanners have yielded large-scale digital collections of high-resolution pathology images. Deep learning models trained on such data are able to classify cell types from images, analyze the tumor microenvironment quantitatively [1, 51], predict tumor subtypes or patient progression, as well as successfully infer molecular features from pathology images [4, 22, 24, 28], including driver mutations and bulk RNA-seq profiles [6, 9, 10]. Such inferences save the effort, cost, and additional specimens required for genomics assays and can be applied to images or archived specimens from cohorts collected over decades and as part of routine clinical care, even in low-resource settings. This then deepens our understanding of how genomics affects morphological phenotypes and can help derive clinically meaningful biomarkers based on combined cellular morphological and molecular phenotypes.

Following the introduction of spatial transcriptomics, researchers have attempted to predict spatial transcriptomics profiles using matched pathology images [8, 31, 49]. However, a significant challenge for such models is the relative lack of paired pathology and spatial transcriptomics data. To address this issue, researchers have actively employed transfer learning strategies that leverage pre-trained pathology image models as an encoder that extracts useful and meaningful embeddings from images. Initially, the effort to leverage models that infer omics profiles from pathology images started with directly fine-tuning pretrained image models that were trained with natural image datasets that do not contain many medical images [42]. However, these were superseded by pathology foundation models that can obtain good representations without annotations by leveraging the ever-increasing size of pathology images and better modeling paradigms [5, 16, 27, 47, 48]. A typical pathology foundation model is trained with over a million whole slide images, and a single pre-trained model surpasses other models that are designed explicitly for downstream tasks such as tumor subtype classification, patient progression prediction, etc [21, 50].

Researchers have now leveraged pre-trained models to obtain high-quality image embeddings from pathology images and use them to infer omics profiles [14, 15, 20, 25]. However, these models have typically used an MLP as an encoder that lacks the capacity to encode omics data effectively into a meaningful embedding space [31, 49]. Recently, more powerful, transferrable models for omics modalities have been introduced. For example, a single-cell foundation model, trained on over 10 million single-cell RNA-seq datasets, has demonstrated performance in tasks that require robust representations, such as single-cell annotation and perturbation prediction [11, 13, 45]. Because spatial transcriptomics profiles are comprised of single-cell profiles organized in space (and possibly partly aggregated), we hypothesized that models can now leverage transfer learning strategies for both imaging modality and spatial transcriptomics modality simultaneously. Here, we introduce PathOmCLIP, a multi-modal pathology-spatial model that leverages powerful foundation models for each modality to predict spatial transcriptomics profiles from pathology images. We aligned the two modalities by applying contrastive loss to embedding vectors extracted from each foundation model, which has shown powerful performance on image-text alignment [35]. While aligning two embedding vectors is beneficial, we found that by itself, it does not adequately capture the multi-cellular context of a given cell, which is key to properly modeling the tumor microenvironment. We thus introduced the LocalTransformer as a mechanism to learn embeddings that are conditioned on their local neighborhood. Integrating LocalTransformer on image embeddings before and after the alignment of the two modalities by contrastive loss significantly improved the prediction of spatial transcriptomics profiles from pathology images. We validated PathOmCLIP using the recently introduced HEST-1K dataset, comparing PathOmCLIP with other competitive models across five different cancer types, including clear cell renal cell carcinoma (ccRCC) [18]. ccRCC is well known for its pathological features that align with patient progression, and we demonstrated that PathOmCLIP effectively infers spatial transcriptomics profile using pathological features of ccRCC [29, 32, 43].

## 2 Related work

### Pathology and Single-cell foundation models

A foundation model is a model that consists of a large number of parameters (e.g., a few million to billions), is trained on large datasets, and can perform a variety of tasks [3, 34]. The model extracts generally representative embedding vectors that can be easily adapted to diverse tasks, even in cases where specific tasks lack sufficient data.

In the pathology domain, there are multiple recent efforts to build foundation models using different modeling approaches and datasets, including multi-modal foundation models that enhance representation power by incorporating both images and their textual descriptions from pathologists. Below, we utilized the GigaPath model [50], which is trained on a dataset that includes diverse tumor types and provides patch-level image features from whole slide pathology images.

Similarly, in single-cell genomics, large-scale curated atlases of single-cell RNA-Seq data, such as the Human Cell Atlas [38, 40] now enabled researchers to develop single-cell foundation models. These are trained on large atlases and can be effectively adapted to a wide range of applications that require robust representation of cells and genes. While some studies have attempted to incorporate spatial transcriptomics data, handling the difference in data characteristics of various spatial omics methods remains somewhat ambiguous [2, 41]. Here, we utilized the scFoundation model [13], which was trained on over 50M single-cell RNA-seq profiles and does not bin the RNA-seq expression profile, making it more suitable for representing spatial omics data.

### Spatial Transcriptomics profile prediction from pathology image

Spatial transcriptomics captures the spatial distribution of RNA profiles across a tissue section. One popular class of approaches uses spotted DNA spatial barcodes to capture the mRNA. With current methods (e.g., Visium, used here) each spot represents the aggregate profile in an area spanning 10-100 cells, and is aligned directly with a corresponding histopathology image stained with hematoxylin and eosin (H&E) at that location. Recently, BLEEP and mclSTExp have been introduced to align these different modalities using contrastive loss [31, 49], and HEST demonstrated the benefits of using pathology foundation models for this task [18]. However, no previous methods have simultaneously used single-cell and pathology foundation models while applying a locally contextualized transformer model to predict spatial expression profiles, which is the approach that we present below.

## 3 Methods

### 3.1 Data and preprocessing

We used the July version of the HEST benchmarking 10x Visium dataset that contains 224×224 H&E stained image patches, each 112×112 μm, and matched spatial gene expression data spot [18]. We included five tumor types: clear cell renal cell carcinoma (kidney cancer, ccRCC), prostate adenocarcinoma (prostate cancer, PRAD), rectal adenocarcinoma (rectum cancer, READ), pancreatic adenocarcinoma (pancreatic cancer, PAAD), colonic adenocarcinoma (colon cancer, COAD) from the HEST dataset. We adhered to their specified training and test splits, which are patient-stratified splits, to avoid patient-level data leakage during the evaluation. We applied log transformation to the raw count of spatial gene expression profile data and unified the gene symbols within the same tumor type, which served as the target spatial gene expression profile.

We used the GigaPath model as our image encoder and followed its image-processing process. Specifically, we normalized each raw 224×224 image from HEST using the mean and standard deviation required by the GigaPath model. We used the scFoundation model for our gene expression encoder and added genes with zero expression to match the input gene symbols required by scFoundation. The log-transformed and count-normalized gene expression data were passed into scFoundation, with the target resolution set to 5 to account for the higher total counts often seen in 10X Visium spatial data compared to single-cell data.

We prepared different numbers of highly variable genes (HVGs) to test the model. We used the 50 HVGs provided by the HEST dataset. For other HVGs, we generated count-normalized and log-transformed gene expression for each tumor type and used the function *scanpy.pp.highly_variable_genes* from the scanpy package and set the *batch_key* indicating the slide to identify HVGs across slides.

### 3.2 PathOmCLIP

PathOmCLIP consists of two-stage training. The first stage is a multi-modal embedding learning through LocalCLIP, and the second stage is training a LocalTransformer for gene expression prediction tasks using the pre-trained LocalCLIP model from stage 1 (**Figure 1**).

**Fig. 1.**
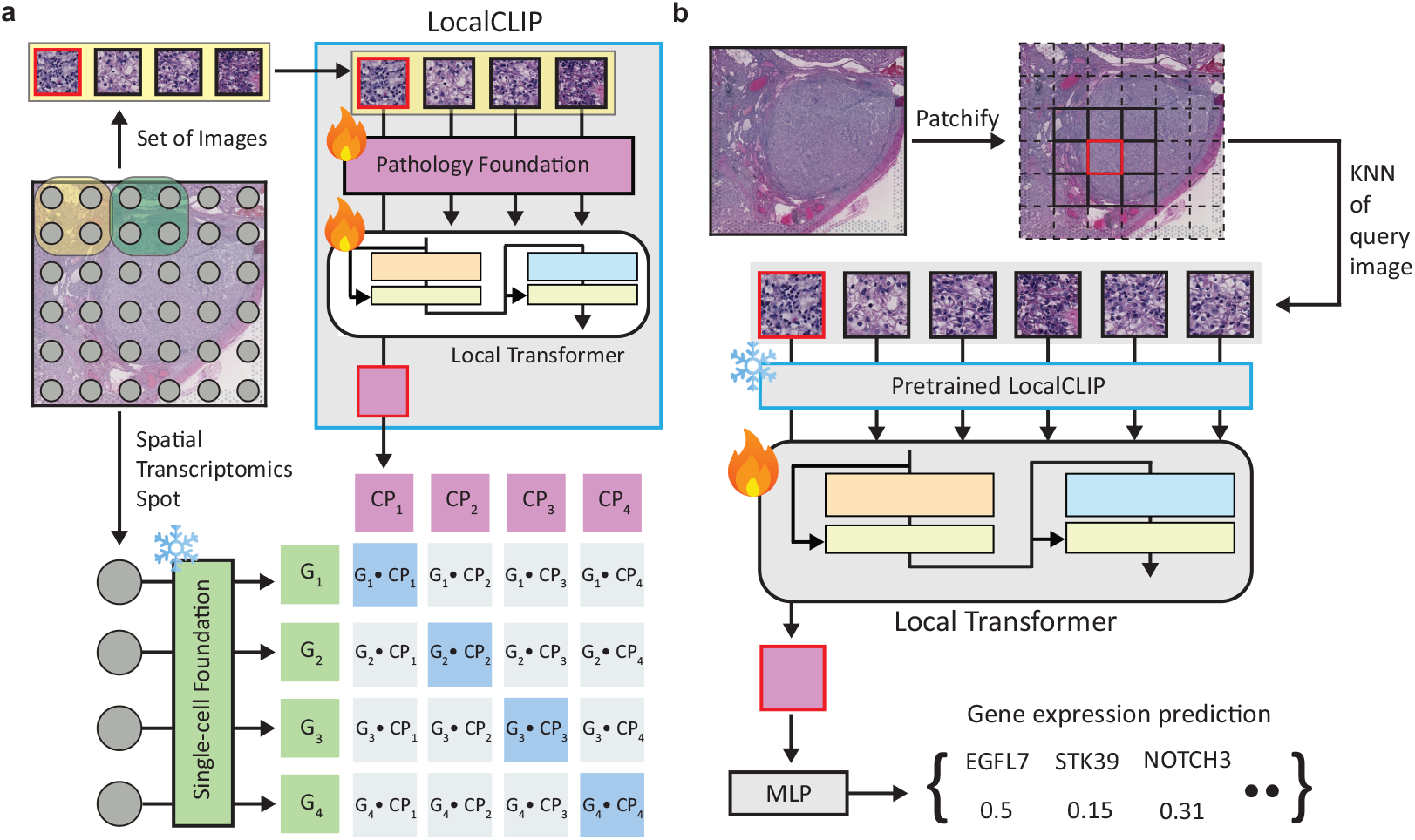
Two-stage training of PathOmCLIP **a**. PathOmCLIP is first trained with a contrastive loss between the embeddings from LocalCLIP and a single-cell foundation model. During the training, only the pathology foundation model and LocalTransformer in LocalCLIP are trainable **b**. PathOmCLIP is further directly trained for spatial gene expression prediction with LocalTransformer. We used the pre-trained LocalCLIP model from stage 1 as inputs to the LocalTransformer and obtained the locally contextualized embedding to predict the gene expression profiles.

#### Learning multi-modal embedding through LocalCLIP

We obtain embeddings from the spatial gene expression profiles of a spot *X* ∈ ℝ^1*×M*^ using the single-cell foundation model as an encoder *f*_*gene*_ : ℝ^*M*^→ ℝ^*h*^that projects the *M* -dim normalized gene expression into a *h*-dim gene expression embedding *H*_*G*_ = *f*_*gene*_(*X*) ∈ ℝ^1*×h*^. We obtain a pathology patch embedding from an image patch *V* ∈ ℝ^1*×C*^ using the pathology foundation model as an encoder *f*_*path*_ : ℝ^*C*^→ ℝ^*d*^ to obtain image embedding *H*_*P*_ = *f*_*path*_(*V*) ∈ ℝ^1*×d*^. The extracted embeddings *H*_*P*_ further go through a LocalTransformer that aggregates local neighborhood information. Let 𝒮_*P*_ = {*H*_*P*_, *H*_1_, *H*_2_, …, *H*_*K*_} be a set of *K* image embedding features provided as input to the Transformer to obtain contextualized image embedding *H*_*CP*_ ∈ ℝ^1*×d*^. 𝒮_*P*_ is updated according to the *k* -nearest neighbor (*k* -NN) image patches of query image V, and we use the patches’ positional coordinates to define the nearest neighbors within the pathology image.

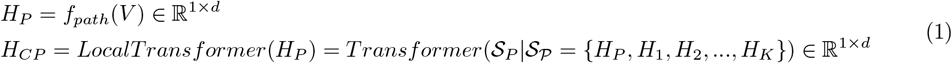

LocalTransformer is a conventional transformer architecture that uses *k* -NN images as input Equation (1). We did not add position embeddings to the input embeddings so that the model is invariant to the position and treat the input image embeddings as a set. We only use the embedding *H*_*CP*_ that corresponds to the contextualized embeddings of *H*_*P*_ and disregard other embeddings from the *k* -NN images’ embeddings. We repeatedly conducted LocalTransformer for *N* images to get the batch-level embeddings.

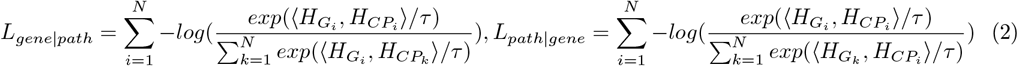

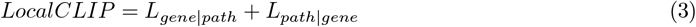

We obtain our gene expression embedding *H*_*G*_ and image embedding *H*_*CP*_ through Equation (1). To align the embeddings from the two modalities, we used contrastive loss. InfoNCE loss showed powerful performance with various multi-modal alignment tasks and specifically visual-text alignment, such as the Contrastive Language-Image Pretraining (CLIP) framework [37]. Unlike image-text alignment, we used infoNCE loss to align the expression and image features, which maximizes the alignment between the true pairs of image and spatial gene expression spot versus others. We maximize the posterior probability of the gene expression embedding of spatial spot *H*_*G*_ given its corresponding image embedding *H*_*CP*_. We also calculated the posterior probability of the image embedding given the expression embedding. The two losses are in Equation (2) where *τ* ∈ R is a temperature parameter and 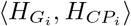 represents the cosine similarity between the matched spatial gene expression embedding and image patch embedding, which combined into our final loss for LocalCLIP training in equation (3).

### Inferring expression profile using LocalTransformer

We obtain the image features *H*_*CP*_ from the pretrained LocalCLIP model. Similar to the LocalCLIP model, we do one additional round of LocalTransformer to use the local neighborhood information directly for the spatial expression prediction task and add one layer of MLP: ℝ^*d*^ → ℝ^*M*^ to infer the gene expression *X*_*pred*_. We use the MSEloss between *X*_*pred*_ and ground truth gene expression *X* to train the model

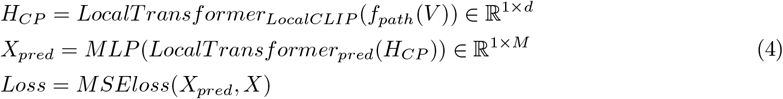

### Training Details

We used the Fabric framework to enable distributed training with bfloat16 precision, running on 4 NVIDIA A100 80GB GPUs. During the stage 1 LocalCLIP training, we set the number of local neighbors to 4, trained for a maximum of 3 epochs, and used a batch size of 64. The learning rate was set to 1e-4 with a weight decay of 1e-5. For the stage 2 LocalTransformer training, we set the number of local neighbors to 50, trained for up to 150 epochs, and used a batch size of 4,096. The learning rate was set to 2e-5 with a weight decay of 1e-5.

### 3.3 Baseline methods

We compared PathOmCLIP with BLEEP and mclSTExp, two established models for predicting spatial expression profiles from pathology images. Both models utilize the alignment of pathology image and spatial gene expression using contrastive loss, and mclSTExp proposed an enhanced spatial expression encoding method before alignment.

Since BLEEP and mclSTExp do not utilize pathology foundation models, we also compared PathOmCLIP with a directly fine-tuned GigaPath model (GigaPath-FT). To fine-tune GigaPath, we fine-tuned the entire tile encoder and employed a linear head for gene expression prediction. We used a batch size of 64, set the learning rate to 1e-4 with a weight decay of 1e-5, and trained for a maximum of 3 epochs.

For BLEEP, to ensure that the numerical value of the Pearson correlation coefficient (PCC) is based on the same expression values, we used the same preprocessed datasets as our model without performing batch removal on both the training and test data. We kept the default model parameters, using ResNet50 as the image encoder, setting the batch size to 256, and training for a maximum of 150 epochs.

For mclSTExp, we used the same datasets and followed their data processing steps to obtain the top 1,000 highly variable genes as the input feature space. We adhered to their default model configuration, setting the batch size to 128, training for a maximum of 90 epochs, and using a 2-layer, 8-head transformer on the spatial gene expression.

## 4 Results

### PathOmCLIP outperforms in predicting spatially resolved expression profiles

We first bench-marked PathOmCLIP on how well the predicted expression profiles correlate with the experimentally measured gene expression compared to BLEEP, mclSTExp, and GigaPath-FT. We measured performance across different numbers of highly variable genes (HVGs) because, although most previous studies have focused on the top 50 HVGs, practical applications require more comprehensive profiles, and the foundation model showed interesting phenomena with a large number of HVGs. PathOmCLIP outperformed all other models across all numbers of HVGs (**Table 1**). Interestingly, GigaPath-FT, the fine-tuned Pathology Foundation model, surpassed the dedicated BLEEP and mclSTExp methods in many cases. However, mclSTExp, which includes multi-modal contrastive training, slightly outperformed GigaPath-FT when a large number of HVGs were considered. This emphasizes the role of contrastive training in aligning pathology image embeddings to spatial gene expression profiles and probably indicates the limitation of predicting expression profiles solely from pathology images.

**Table 1.** Average Pearson Coefficient Correlation (PCC) of predicted spatial gene expression compared to ground truth expressions across different numbers of highly variable genes (HVGs). We used the ccRCC dataset that includes ten slides for testing (GigaPath-FT: Finetuned GigaPath)

| Methods | 50 HVGs | 200 HVGs | 500 HVGs | 1000 HVGs | 2000 HVGs |
| --- | --- | --- | --- | --- | --- |
| BLEEP | 0.116±0.055 | 0.079±0.053 | 0.064±0.037 | 0.052±0.025 | 0.051±0.023 |
| mclSTExp | 0.190±0.079 | 0.134±0.052 | 0.118±0.037 | 0.098±0.027 | 0.101±0.032 |
| GigaPath-FT | 0.219±0.040 | 0.154±0.052 | 0.128±0.052 | 0.103±0.031 | 0.098±0.023 |
| PathOmCLIP | **0.288±0.036** | **0.205±0.035** | **0.172±0.026** | **0.133±0.021** | **0.128±0.025** |

### PathOmCLIP predicted gene expression profiles maintain spatial expression pattern

Although a high PCC could indicate good inference performance, another important measure is whether the spatial relationship between the cells in the tissue is maintained. To assess this, we clustered the spot profiles by Leiden clustering with different cluster numbers and evaluated how similar the cluster assignment for each spot is between the measured and predicted expression profiles based on the adjusted mutual information (AMI) (**Supplementary Materials**). PathOmCLIP outperformed all methods with the highest AMI scores (**Figure. 2a**) and a striking similarity to the spatial pattern of the ground truth compared to GigaPath-FT and mclSTEx (**Figure. 2b**). Thus, while the GiGaPath-FT model’s PCC score is high (albeit lower than PathOmCLIP), it does not necessarily preserve the spatial relationship between cells in the tissue (**Figure. 2b**). The critical difference between GigaPath-FT, mclSTExp, and PathOmCLIP is the LocalTransformer module that is applied to the image embeddings. mclSTexp has a similar module that enforces the encoding of neighborhood information, but it is applied to expression profiles rather than to image embeddings. We thus conclude that using the LocalTransformer for image embedding is crucial for achieving both high PCC and well-maintained spatial information.

**Fig. 2.**
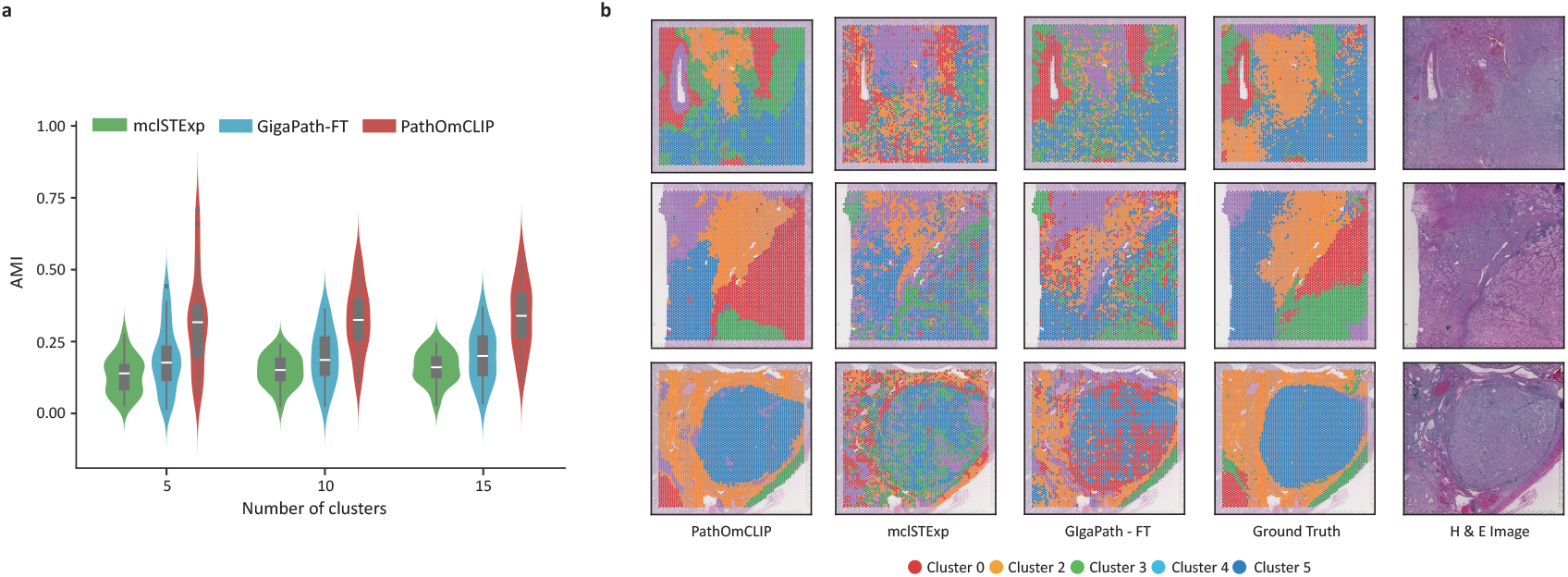
Clustering results of measured (ground truth) and predicted expression of different models overlaid on the pathology image. Each method runs over ten ccRCC slides, and the error bar is the 95% CI of the mean. **a**. AMI score (y-axis) of three different models with three different cluster numbers to create the clustering. **b**. Leiden cluster assignment of each spot (color) overlaid on the pathology image.

### Pathology foundation model is insufficient for predicting spatial gene expression profiles

As noted above, fine-tuning the pathology foundation model surpassed other models but not PathOmCLIP, indicating that the pathology foundation model alone is insufficient to achieve the highest performance (**Table. 1**). Importantly, genes with significant differences between GigaPath-FT and PathOmCLIP predictions in ccRCC (**Figure. 3a**) include key genes with important roles in tumors, such as those related to cell proliferation (LY6E, STK39, NOTCH3) or angiogenesis (EGFL7, HTRA1) [30]. For example, GigaPath-FT predicts that EGFL7 and STK39 are expressed almost uniformly across the tissue, whereas PathOmCLIP distinguishes their expression across space more consistently with the measured ground truth (Figure. 3b).

**Fig. 3.**
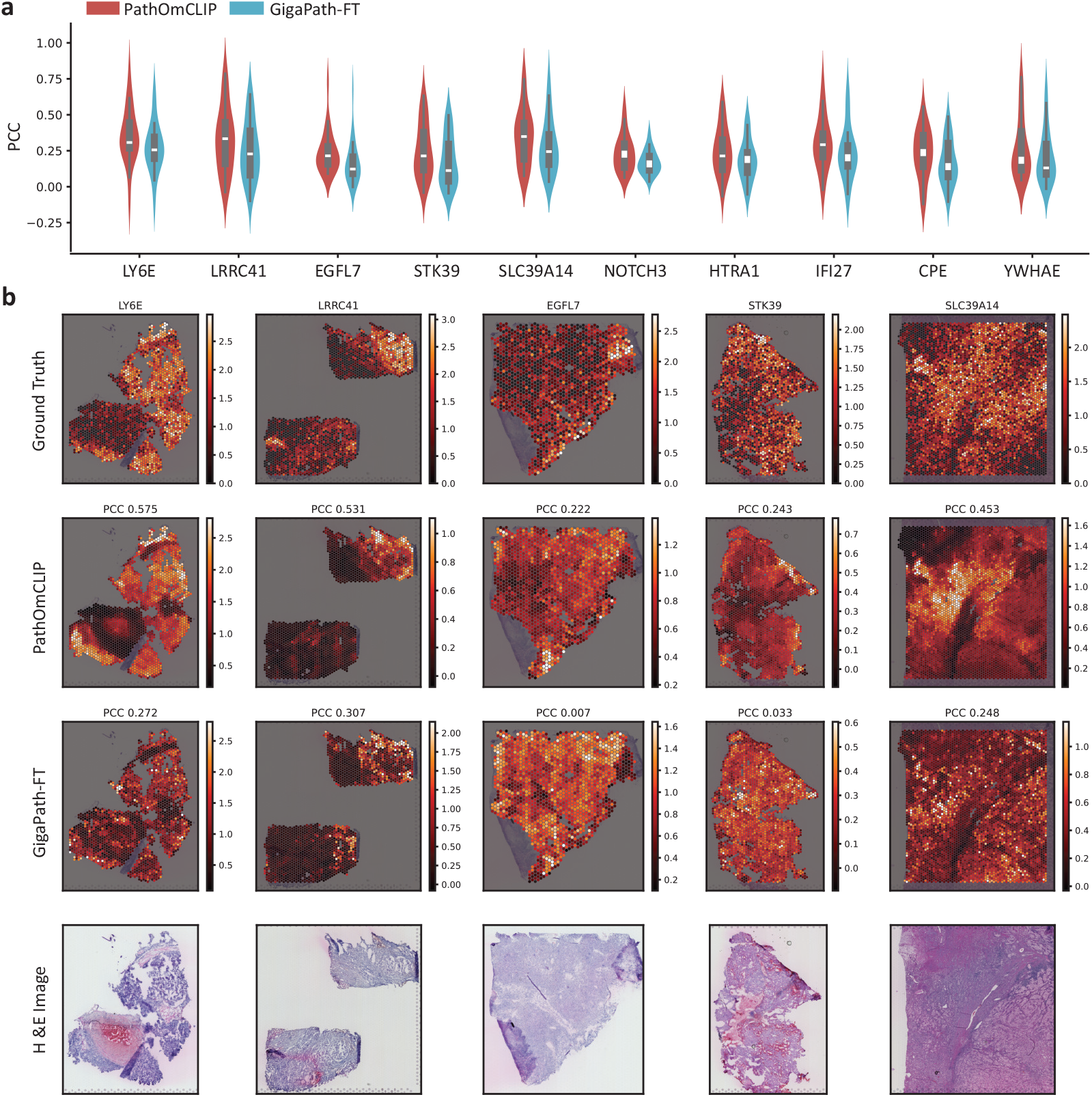
Difference between GigaPath-FT and PathOmCLIP model. **a**. Selected the top ten genes that showed the high PCC difference between the two models. We measured it across ten ccRCC slides, and the error bar is the 95% CI of the mean. **b**. Visualization of ground truth and predicted gene expression profiles. Color map indicates the log-transformed gene expression profiles

### Ablation studies highlight the importance of LocalCLIP and a single-cell foundation model

We next conducted ablation studies to understand which design components are critical for PathOmCLIP performance (**Table. 2**). Adding LocalTransformer slightly increases the performance compared to the naive GigaPath model (Ridge Regression). Adding CLIP loss on top of the LocalTransformer boosts the performance significantly, and when we tested the model on 50 HVGs, performance seemed to drop when we used the single-cell foundation model (scFoundation) or LocalCLIP, which is the critical component of PathOmCLIP. Interestingly, applying LocalCLIP on both modalities harms the performance, which shows the difference between the modalities. We speculate that the pathology image has a higher local similarity compared to spatial gene expression, so guiding locally enhanced features on the image is helpful for performance. Conversely, when we increased the number of target genes beyond 50 HVGs, the benefit of the single-cell foundation model and LocalCLIP was apparent (**Table. 3**). This suggests that MLP can overfit to 50 HVGs, which makes the scFoundation model seem less powerful than an MLP.

**Table 2.** Ablation study includes different contrastive loss and gene expression encoders. Average PCC on five-fold cross-validation ccRCC test sets. We tested the methods for 50 HVGs. (LT: *LocalTransformer*_*pred*_, BLocalCLIP: LocalCLIP applied for both image and spatial gene expression for CLIP training)

| Methods | Split 0 | Split 1 | Split 2 | Split 3 | Split 4 | Mean |
| --- | --- | --- | --- | --- | --- | --- |
| LT | 0.131 | 0.235 | 0.279 | 0.193 | 0.277 | 0.223±0.06 |
| Ridge Regression | 0.176 | 0.298 | 0.166 | 0.228 | 0.221 | 0.218±0.05 |
| LT + CLIP + scF | 0.245 | 0.274 | 0.270 | 0.241 | 0.312 | 0.268±0.02 |
| LT + CLIP + MLP | 0.269 | 0.345 | 0.316 | 0.235 | 0.286 | **0.290±0.04** |
| LT + LocalCLIP + scF | 0.239 | 0.296 | 0.298 | 0.337 | 0.271 | <u>0.288±0.03</u> |
| LT + LocalCLIP + MLP | 0.236 | 0.281 | 0.208 | 0.375 | 0.290 | 0.278±0.06 |
| LT + BLocalCLIP + scF | 0.251 | 0.256 | 0.288 | 0.301 | 0.278 | 0.275±0.02 |
| LT + BLocalCLIP + MLP | 0.226 | 0.317 | 0.189 | 0.292 | 0.297 | 0.264±0.05 |

**Table 3.** Ablation study tests the effect of LocalTransformer for contrastive loss and a single-cell foundation model as a spatial gene expression encoder. MLP indicates the one-layer MLP for spatial gene expression encoders. Average PCC of ten ccRCC test slides. Tested across 50, 200, 500, 1000, and 2000 HVGs. (*∗* is the PathOmCLIP, scF: Single-cell foundation model)

| Methods | 50 HVGs | 200 HVGs | 500 HVGs | 1000 HVGs | 2000 HVGs |
| --- | --- | --- | --- | --- | --- |
| GigaPath-FT | 0.219±0.040 | 0.154±0.052 | 0.128±0.052 | 0.103±0.031 | 0.098±0.023 |
| CLIP w/ MLP | **0.290±0.042** | 0.179±0.059 | 0.139±0.062 | 0.118±0.037 | 0.112±0.053 |
| CLIP w/ scF | 0.268±0.029 | 0.185±0.061 | <u>0.156±0.043</u> | 0.126±0.022 | <u>0.121±0.030</u> |
| LocalCLIP w/ MLP | 0.278±0.064 | <u>0.192±0.053</u> | 0.155±0.037 | <u>0.127±0.016</u> | 0.120±0.025 |
| *LocalCLIP w/ scF | <u>0.288±0.036</u> | **0.205±0.035** | **0.172±0.026** | **0.133±0.021** | **0.128±0.025** |

Adding CLIP loss to the pathology foundation model increased performance compared with the GigaPath-FT model, even without the scFoundation model. However, if we increase the number of genes for prediction to 200 HVGs, using the scFoundation model as an encoder shows better performance compared to MLP. Introducing LocalCLIP shows a similar pattern of impact in that it harms performance on the top 50 HVGs and increases performance on the top 200 HVGs. Finally, when we introduce LocalCLIP with scFoundation, it always shows better performance compared to using MLP as a spatial omics encoder, which supports our design choice to add LocalTransformer before and after CLIP training and using scFoundation as the encoder.

### PathOmCLIP leverages multi-modal information to refine the representation

We further tested whether a single-cell foundation model provides advantages across different tumor types. Similar to ccRCC, incorporating a single-cell foundation model was beneficial when the number of target HVGs increased across four other tumor types (**Table. 4**). We also examined the effect of multi-modal training on the embedding space. To this end, we visualized the image features obtained from PathOmCLIP and GigaPath. To explore the effect of each training step and component, we visualized features after stage 2 LocalTransformer, stage 1 LocalCLIP, and stage 1 LocalCLIP but using MLP as a spatial gene expression encoder. In addition, we visualized the spatial gene expression features from scFoundation (**Figure. 4a**). We measured the average silhouette width (ASW) score of each embedding space, and we used each slide label as a cluster label, which means a lower ASW score indicates the features obtained from each slide mixed better. CLIP training resulted in a clearer mixing of certain embeddings through multi-modal information, which cannot be achieved by either single-cell-only embedding or pathology embedding alone. The PathOmCLIP that used scFoundation as an omics encoder shows lower ASW than the MLP one, demonstrating the benefit of using a pre-trained single-cell foundation model. Stage 2 LocalTransformer mitigates the batch effect between each slide better than LocalCLIP by making the embedding vectors include more contextualized information. Overall, this demonstrates the power of the multi-modal training and foundation model. We also confirmed the above strong performance by visualizing the predicted spatial gene expression compared with previous models. We visualized the difference between the single-cell foundation model and a simple MLP encoder. In the case of TRBC1 and HSPA1A, scFoundation clearly helps to reduce the false-positive prediction (**Figure. 4b**).

**Table 4.** PathOmCLIP’s spatial gene expression inferring performance across different cancer types. Average PCC of each cancer type that tested across 50, 200, 1000, and 2000 HVGs. Over 1000 HVGs of PAAD is unavailable because of not enough captured genes

| Cancer Type | Methods | 50 HVGs | 200 HVGs | 1000 HVGs | 2000 HVG |
| --- | --- | --- | --- | --- | --- |
| PRAD | PathOmCLIP w/ MLP | 0.330±0.017 | 0.134±0.010 | 0.109±0.024 | 0.106±0.026 |
|  | PathOmCLIP w/ scF | **0.381±0.014** | **0.140±0.011** | **0.121±0.013** | **0.120±0.016** |
| READ | PathOmCLIP w/ MLP | 0.199±0.069 | 0.121±0.059 | **0.045±0.010** | 0.019±0.003 |
|  | PathOmCLIP w/ scF | 0.199±0.081 | **0.125±0.056** | 0.043±0.014 | **0.020±0.008** |
| PAAD | PathOmCLIP w/ MLP | 0.402±0.014 | 0.268±0.012 | N\A | N\A |
|  | PathOmCLIP w/ scF | **0.464±0.016** | **0.317±0.026** | N\A | N\A |
| COAD | PathOmCLIP w/ MLP | **0.200±0.059** | **0.181±0.034** | 0.108±0.025 | 0.067±0.017 |
|  | PathOmCLIP w/ scF | 0.164±0.020 | 0.169±0.034 | **0.116±0.027** | **0.080±0.017** |

**Fig. 4.**
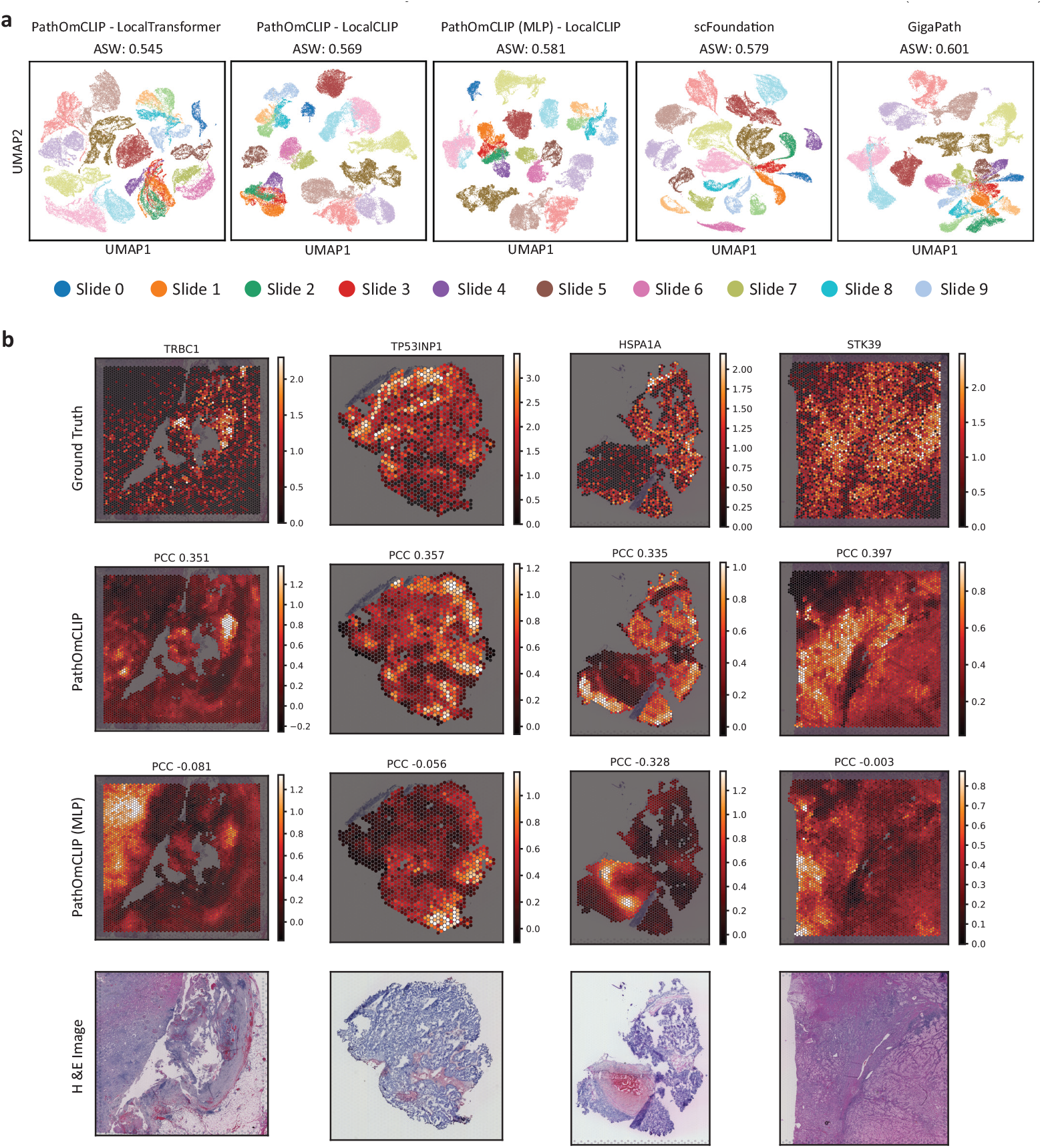
Benefit of scFoundation over MLP as gene expression encoder. **a**. UMAP plot of features obtained from ten ccRCC slides. The features were collected after stage 2 LocalTransformer, stage 1 LocalCLIP of the PathOmCLIP model, and LocalCLIP of the PathOmCLIP with MLP as the gene expression encoder. Additionally, features were collected from scFoundation and GigaPath. The ASW scores were calculated using each slide label as a cluster label. **b**. Visualization of ground truth and predicted gene expression profiles that showed the large difference between PathOmCLIP trained with scFoundation and MLP. Color map indicates the log-transformed gene expression profiles

## 5 Discussion

Our study demonstrates that leveraging a foundation model of each modality, specifically a single-cell foundation model and a pathology foundation model, enhances the prediction accuracy of spatial transcriptomics profiles from pathology images. In addition, integrating local information is critical to achieving the highest performance with the alignment of multi-modal information. Although PathOmCLIP demonstrates the benefit of using foundation models for both modalities, it still has some limitations. First, we tested the combination of scFoundation as a spatial gene expression encoder and GigaPath as a pathology image encoder, but there are recently introduced more advanced single-cell foundation models, spatial transcriptomics models, and pathology foundation models as well [2, 41]. Different pairs of these foundation models may further affect the overall performance, and finding the optimal pair should be further explored. Second, batch effects in each modality further impact performance. Recent studies have shown that pathology foundation models can handle batch effects within samples or patients but still suffer from batch effects between patients, and single-cell foundation models cannot easily handle this problem [23, 26, 52]. When conducting multi-modal training, batch effects can be mitigated, but they still affect the overall training performance, so we need a better solution for the multi-modal batch effect problem. Transfer learning helps to boost performance, but the model requires more paired data to test its reliability. HEST-1K is the largest curated spatial transcriptomics data, but some tumor types only have samples from as few as two patients. Additionally, most pathology foundation models are trained with pathology images scanned from FFPE samples, but many samples collected for spatial transcriptomics are fresh-frozen samples, which induce the performance drop when directly transferring the model trained mainly on the FFPE sample. Recently introduced generative models that convert fresh-frozen into FFPE images or spatial transcriptomics methods that can collect omics data from FFPE samples can help address this problem [36]. Finally, we need further interpretation of how these models predict the gene expression profile using morphological features. Higher PCC performance compared with ground truth measured expression does not always guarantee that the model learns meaningful features. Understanding the connection between morphology and genomics using these models in collaboration with pathologists could be an interesting topic to explore.

Overall, inferring molecular features from pathology images could be clinically beneficial when we apply this model to many archived histology images, unlocking valuable clinical information and facilitating new biomarker discoveries.

## Acknowledgements and Funding Information

We thank Wan Changlin and Hector Corrada Bravo for their feedback on how to improve the manuscript. We are grateful to the members of the Regev Lab for their valuable feedback on earlier drafts of the results. AR is an employee of Genentech and has equity in Roche, and is a co-founder and equity holder of Celsius Therapeutics and an equity holder in Immunitas. She was an SAB member of Thermo Fisher Scientific, Syros Pharmaceuticals, Neogene Therapeutics, and Asimov until July 31st, 2020.

## Code Availability

The raw source code is available upon request.

## 6 Supplementary

### Clustering analysis

We performed clustering analysis based on the 2,000 HVG predictions of the top two methods: ours and mclSTExp. We used the log-transformed raw gene expression data as the ground truth. For each slide, we conducted PCA and constructed a neighbor graph with default parameters. Clustering was performed with 5, 10, and 15 clusters per dataset. To ensure consistency in the number of clusters, we employed a modified Leiden algorithm that dynamically adjusts resolution to match the desired cluster count. We then evaluated performance using the adjusted mutual information (AMI) score between the predicted clusters and the ground truth. For visualization, we applied the Hungarian algorithm to align cluster colors across methods.

### Overall Model Structure

We use *GigaPath* for our image encoder and *scFoundation* as our gene expression encoder. The 4-layer transformers with a single attention head are used for *LocalTransformer*_*LocalCLIP*_. To match the dimensions of the two modalities, we use a linear image projector and a linear expression projector. The image encoder outputs a sequence of 1,536-dimensional embeddings, which the linear image projector reduces to 512 dimensions. The expression projector similarly encodes expression data into a 512-dimensional space, and the CLIP loss is computed between the encoded expression embeddings and the local image embeddings. Our *LocalTransformer*_*pred*_ model features a 4-layer transformer with one attention head and a linear prediction head. The prediction head processes the first embedding from the transformer’s output to generate a gene expression vector.

